# Protein Thermal Stability Engineering Using Hotmusic

**DOI:** 10.1101/539528

**Authors:** Fabrizio Pucci, Jean Marc Kwasigroch, Marianne Rooman

## Abstract

The rational design of enzymes is a challenging research field, which plays an important role in the optimization of a wide series of biotechnological processes. Computational approaches allow to screen all possible amino acid substitutions in a target protein and to identify a subset likely to have the desired properties. They can thus be used to guide and restrict the huge, time-consuming, search in sequence space to reach protein optimality. Here we present HoTMuSiC, a tool that predicts the impact of point mutations on the protein melting temperature, which uses the experimental or modelled protein structure as sole input, and is available at dezyme.com. Its main advantages include accuracy and speed, which makes it a perfect instrument for thermal stability engineering projects aiming to design new proteins that feature increased heat resistance or remain active and stable in non-physiological conditions. We set up a HoTMuSiC-based pipeline, which uses additional information to avoid mutations of functionally important residues, identified as being too well conserved among homologous proteins or too close to annotated functional sites. The efficiency of this pipeline is successfully demonstrated on *Rhizomucor miehei* lipase.

## 1. Introduction

In the last decades, lots of efforts have been devoted to analyse the molecular mechanisms associated to protein thermal stability, since their understanding is fundamental not only for advancing the theoretical comprehension of the protein folding process, but also for potential applications to a wide series of biological processes that range from drug design to the synthesis of new protein nanomaterials. The design of new enzymes that remain active and stable at temperatures well above or below their physiological temperature can also lead to the improvement of the efficiency of catalytic processes, while reducing their economic costs and their environmental impact [1–3]. Different experimental and computational approaches have been developed and largely used to enhance protein thermal resistance [4–6] such as:

- Directed evolution methods in which randomly distributed mutations are introduced in the target protein and are followed by screening and selection steps.
- Rational protein design in which the understanding of the protein structure/function relationships is the key ingredient.
- Semi-rational approaches that combine the benefits of the directed evolution and the rational design methods.

Despite impressive current achievements, the development of a systematic and cheap way to optimize a target protein remains a challenging goal. This is primarily due to the huge size of the sequence space to be explored, and to the incomplete knowledge of the molecular mechanisms involved. Here we present some advances in the computational protein design field by describing the HoTMuSiC software [7] that we recently developed. HoTMuSiC is an efficient tool that, given the experimental or modelled three-dimensional (3D) structure of the target protein as input, screens the protein sequence and predicts the impact of all possible single amino acid substitutions on the protein melting temperature in just a few minutes. Its performance makes it a perfect instrument to design proteins with improved thermal stability and to guide mutagenesis experiments that are usually expensive and time-consuming. Moreover, due to its speed, it can also be employed in large-scale investigations aimed to gain insights into the relationship between protein thermal stability and natural evolution, and into the adaptation mechanisms to extreme environmental conditions [8]. In the next sections, we briefly introduce the key ingredients and model structure of HoTMuSiC, show how it can be fruitfully applied for protein design applications, and discuss its performances in detail.

## 2. HoTMuSiC key instrument: the statistical potentials

The key instrument used in the construction of HoTMuSiC is the statistical potential formalism, known to be quite efficient when applied to a wide range of problems including protein structure prediction, protein design and protein-ligand scoring. These knowledge-based, effective, potentials are derived from frequencies of sequence and structure elements in a non-redundant dataset of well resolved protein 3D structures [9]. More precisely, let c be a structure element, consisting of the distance between two residues, the solvent accessibility of a residue, its backbone torsion angle, or combinations thereof, and let s be a sequence element consisting of the amino-acid type of one or two residues. The free energy contribution of the association (c, s) is ruled by the Boltzmann law:

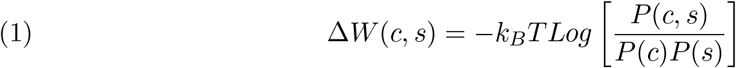

where *P*(*c*), *P*(*s*) and *P*(*c, s*) are the relative frequencies of *c*, *s* and (*c*, *s*) in the dataset, *k_B_* is the Boltzmann constant and *T* the absolute temperature. In order to construct energy functions that describe the impact of the temperature on the amino acid interactions, we introduced temperature-dependent statistical potentials [10, 11]. They are extracted from datasets of proteins with known structures and specified thermal stability properties. The potentials derived from mesostable proteins, denoted as Δ*W*(*c, s*)*^meso^*, describe the interactions at low T, while those derived from thermostable or hyperthermostable proteins, Δ*W*(*c, s*)*^thermo^*, represent the interactions at high temperatures. Using these new potentials, the T-dependence of several types of amino acid interactions were unravelled [10]. Moreover, these potentials were successfully applied to the prediction of the protein stability curve as a function of the temperature [11, 12]. The potentials that are used in HoTMuSiC’s model are the standard potentials Δ*W*(*c, s*) and the T-dependent potentials Δ*W*(*c, s*)*^meso^* and Δ*W*(*c, s*)^thermo^, for various structure and sequence elements (*c, s*).

## 3. HoTMuSiC Harmony: the artificial neural networks

The statistical potentials are combined to predict how the melting temperature Tm changes upon amino acid substitution:

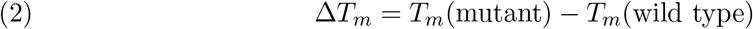

We set up two different models to compute Δ*T_m_*. The HoTMuSiC model requires as sole input the 3D structure of the wild type protein and is based on a linear combination of the standard statistical potentials Δ*W*(*c, s*) for different sequence and structure elements (*c, s*). The Tm-HoTMuSiC model requires in addition the melting temperature of the wild type protein and employs a combination of standard and T-dependent potentials Δ*W*(*c, s*), Δ*W*(*c, s*)*^thermo^* and Δ*W*(*c, s*)*^meso^*. Both models include three additional terms: a constant term 1, and two terms Δ*V*± that represent the difference in volume between the wild-type and mutant residues, and describe the creation of stress or holes in the protein structure. All these terms are weighted by sigmoid functions of the solvent accessibility A, which smoothly connect the surface to the protein interior, while allowing different behaviours in these regions. Indeed, according to the type of potential, the mutational impact can be stronger at the surface or in the protein core. Each sigmoid is a function of four parameters, which were identified using as cost function the difference between experimental and computed Δ*T_m_* values on a learning dataset. We used for that purpose the T1626 set [13] that contains 1,626 mutations inserted in about 90 proteins of known X-ray structure of resolution below 2.5 Å, collected by literature screening. For minimizing the cost function, artificial neural networks (ANN) were considered. For HoTMuSiC, a standard single layer ANN was used (Fig. 1a), in which each perceptron is associated with one input term (Δ*W*(*c, s*), Δ*V*±, 1), and the activation functions are the sigmoid functions of *A*. The ANN of *T_m_*-HoTMuSiC is composed of three layers, where the additional - hidden - layer gets activated by functions of the *T_m_* of the wild-type protein, and confer more weight to mesostable or thermostable perceptrons according to the thermal properties of the target protein (see Fig. 1b and [7] for details). A standard back-propagation algorithm was employed in the training of the neural network. To test the algorithm, we performed 5-fold cross validation using an early stopping technique to avoid overfitting.

**Figure 1.**
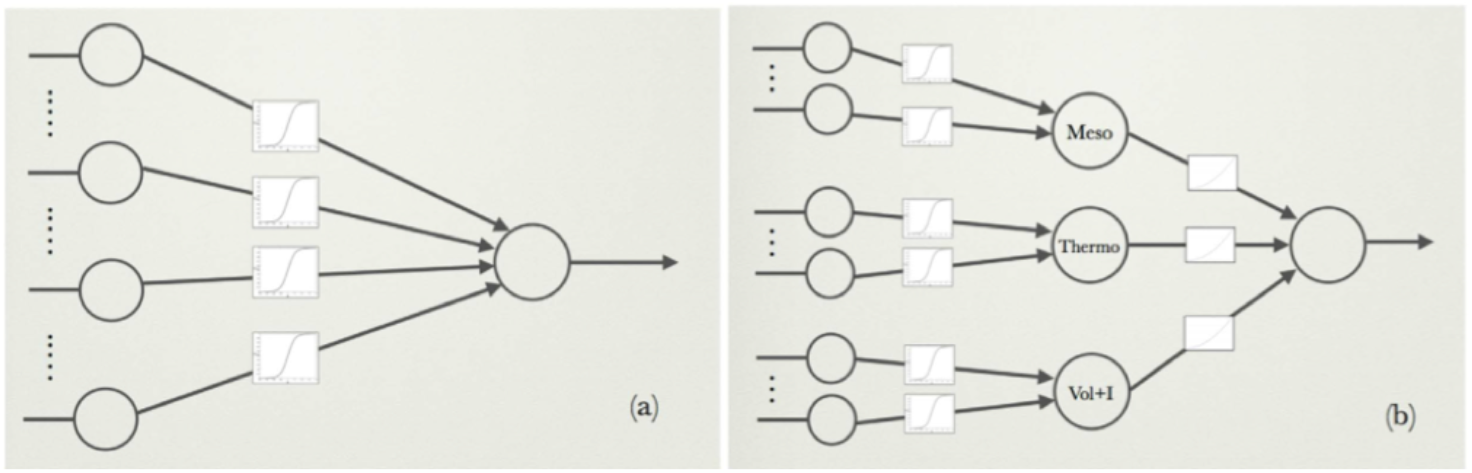
Schematic representation of the ANNs used for the parameter identifications. (a) HoTMuSiC: 2-layer ANN, consisting of a perceptron with sigmoid activation functions and 12 input neurons encoding 9 potentials, two volume terms and a constant term; (b) *T_m_*-HoTMuSiC: 3-layer ANN, consisting of 3 perceptrons with sigmoid weights; the first two perceptrons have 5 input neurons each, encoding 5 mesostable and 5 thermostable potentials, respectively; the third perceptron has 3 neurons for the volume and constant terms. The outputs of these three perceptrons (Meso, Thermo, and Vol + I) are the inputs of another perceptron with polynomial weight functions of the wild-type Tm value.

## 4. HoTMuSiC sound: the results

### 4.1. Performances

To validate the method, we first applied the two HoTMuSiC models to the T1626 learning dataset using a 5-fold cross validation procedure, and compared their performances with another Δ*T_m_* predictor developed in the literature, i.e. AutoMute [14]. The results are reported in [7]. Here, in addition, we compared the HotMuSiC performances with popular ΔΔ*G* prediction methods, namely PoPMuSiC [15], FoldX [16] and Rosetta [17]. Indeed, there is the tendency in the literature to use thermodynamic stability prediction methods to compute thermal stability changes even though the two quantities ΔΔ*G* and Δ*T_m_* are only partially correlated [13]. These comparisons give us an idea of the additional error that one makes by using ΔΔ*G* predictors to predict Δ*T_m_*. The results are given in Table 1.

**Table 1.**
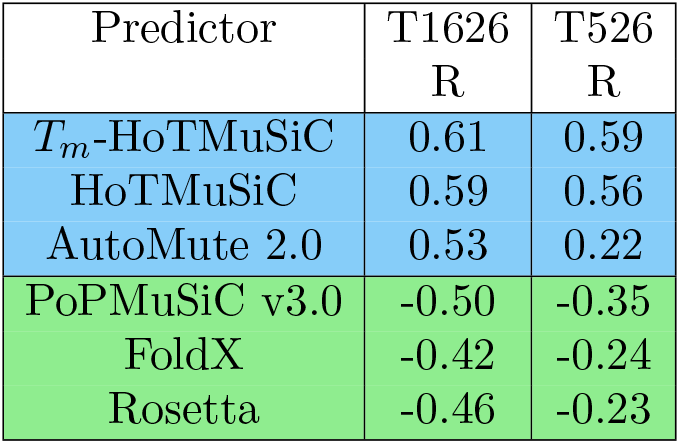
Performances of the predictors on the T1626 [13] and T526 datasets as evaluated by the Pearson correlation coefficients R between the predictor output and the experimental Δ*T_m_* values. The three predictors on a blue background are Δ*T_m_* predictors and the three on a green background are ΔΔ*G* predictors; the latter show negative correlations as, by convention, negative ΔΔ*G* and positive Δ*T_m_* values are stabilizing.

| Predictor | T1626<br>R | T526<br>R |
| --- | --- | --- |
| $T_m$-HoTMuSiC | 0.61 | 0.59 |
| HoTMuSiC | 0.59 | 0.56 |
| AutoMute 2.0 | 0.53 | 0.22 |
| PoPMuSiC v3.0 | -0.50 | -0.35 |
| FoldX | -0.42 | -0.24 |
| Rosetta | -0.46 | -0.23 |

To extend the performance analysis and get more robust results, we also did a manual literature search and used dedicated annotation servers such as ProTherm [18] and Brenda [19] to collect mutations that are not in T1626, i.e. newly characterized substitutions inserted in experimentally solved protein structures, and mutations inserted in structures that are not yet solved but were obtained by homology modelling. In this way, we obtained a second dataset, called T526, containing 526 mutations in 42 experimental and 58 modelled protein structures.

As seen in Table 1, *T_m_*-HoTMuSiC is slightly more accurate than HoTMuSiC, which is normal as it relies on additional information, i.e. the experimental *T_m_* of the wild type protein. Both HoTMuSiC models outperform the other Δ*T_m_*-predictor, especially on the T526 test set, which is not part of any of the predictors’ learning sets. This could indicate that this predictor suffers from overfitting problems.

Moreover, the performance of ΔΔ*G* predictors is significantly lower than that of HoTMuSiC on both sets T1626 and T526. We may thus conclude that using ΔΔ*G* predictors to predict Δ*T_m_* leads to a drop of the performance, of at least 0.1 in correlation coefficient.

### 4.2. Webserver

HoTMuSiC and *T_m_*-HoTMuSiC are freely available for academic use on the webserver dezyme.com. The site contains an introductory and an explanatory page (**Software** and **Help**), and three working pages:

- **My Data** privately stores the user’s personal files, containing protein structures in PDB [20] format or lists of mutations.
- **Query**: on the “HoT” query page, the user must either choose a PDB code that is automatically retrieved from the PDB data bank [20] or a private structure file stored in his My Data page. He must also choose among three options:

1. Systematic mutation option in which all the possible amino acid substitutions for each residue in the protein sequence are computed.
2. Uploading a mutation File containing a list of single-site mutations.
3. Giving Manually a list of single-site mutations. The last option gives the choice between HoTMuSiC and *T_m_*-HoTMuSiC. When selecting the latter, the *T_m_* of the wild type must be specified.
- **The Results** page contains all the predictions performed by the user, which can easily be downloaded. If the systematic mutation mode is chosen, the following files are available:

1. (pdbname).hot contains the predicted Δ*T_m_* value for all possible single-site substitutions inserted in the protein of structure (pdbname) as well as other biophysical characteristics of the mutated residue, i.e. its solvent accessibility and its secondary structure.
2. (pdbname).hots contains information at the residue level: the mean Δ*T_m_* of all substitutions at each position, and the sum of all positive and of all negative Δ*T_m_* values.
3. (pdbname).html shows a histogram picture in which the sum of the Δ*T_m_* of all the stabilizing mutations at each position is indicated. Fig. 2 contains an example of this output.
4. (pdbname.zip) is an archive containing the three abovementioned files, ready to download.

**Figure 2.**
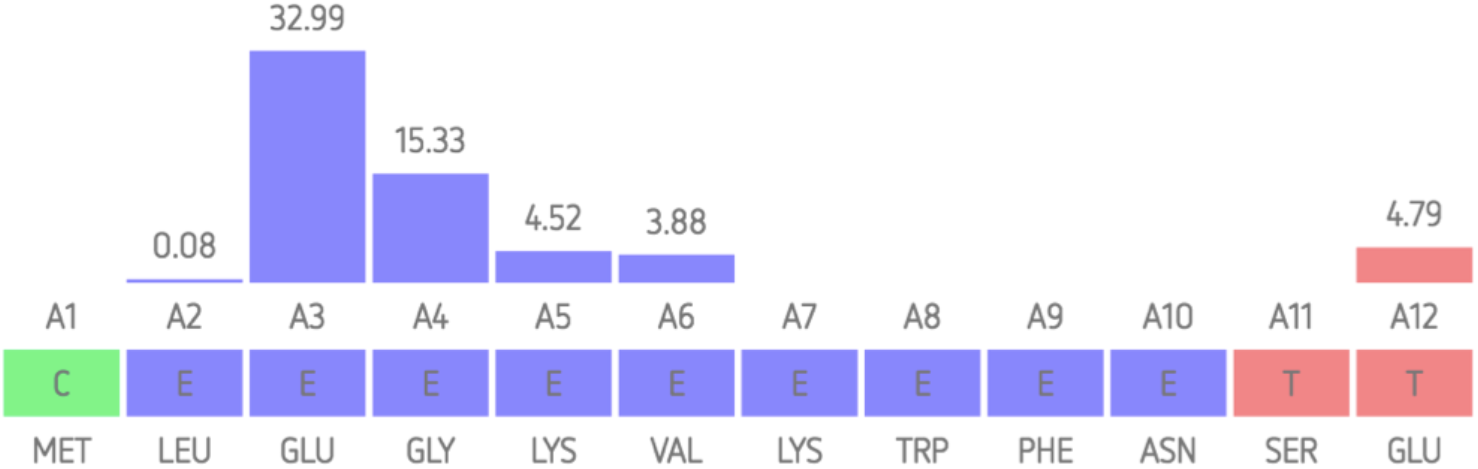
Example of the webserver’s histogram results for the systematic scanning of a target protein. The bars represent the sum of the positively predicted Δ*T_m_* values at each sequence position; these sums are indicated above the bars (in °C). The largest bars represent residues whose mutation are likely to thermostabilize the protein and on which protein engineering experiments should focus first.

### 4.3. Application to modelled structures

The quality of the input structures is of fundamental importance for stability predictions. Indeed, the more precise the structure in terms of resolution, the more likely the predictions are to be accurate. This point is often overlooked when the performances of predictors are discussed. However, it will probably become even more important in the near future since, for example, a growing number of structures will be resolved with cryo-electron microscopy techniques, which usually have lower resolutions (at least for now) than those obtained with X-ray crystallography.

In the newly developed dataset T526, we have also included structures obtained via homology modelling using the Swiss-Model server [21], in view of analysing the robustness of HoTMuSiC with respect to structural inaccuracies. We compared the performance on proteins for which we have a good-resolution X-ray structure, or a modelled structure obtained with a template of which the sequence identity (SI) with respect to the target is either greater than 98% or between 22% and 98%.

As seen from Table 2, the proteins whose X-ray structure is available or can reliably be modelled from templates with SI≥ 98% show good performances with a linear correlation coefficient of about 0.60, whereas the proteins that are modelled from templates SI <98% (with a mean 〈*SI*〉 of 69%) have, as expected, lower but still reasonably good performances measured by a correlation coefficient R≈ 0.45.

**Table 2.** Performances of the predictors on the T526 datasets as evaluated by the Pearson correlation coefficients R between the predictor output and the experimental Δ*T_m_* values. “≥98%” and “≤98%” indicate the sequence identity with respect to the template used for homology modelling.

| Predictor | R |  |  |
| --- | --- | --- | --- |
| | X-ray | $\geq 98\%$ | $\leq 98\%$ |
| $T_m$-HoTMuSiC | 0.59 | 0.62 | 0.45 |
| HoTMuSiC | 0.56 | 0.58 | 0.46 |

Another important aspect that impacts the prediction accuracy is the experimental technique used to determine protein structures and the associated resolution. Clearly, the best results are obtained with X-ray structures of resolution of 2.0-2.5 Å at most, as expected from the above analysis on modelled structures.

These results show that HoTMuSiC can be reliably applied not only to good-resolution structures, but also to low-resolution and modelled structures. This further broadens the field of applicability of this method.

### 4.4. Protein design with HoTMuSiC pipeline

In order to optimize the thermal stability of a target protein to allow it, for example, to remain stable and active at temperatures higher than the physiological temperature, one has to select mutations that increase the protein melting temperature without affecting the protein function. We constructed a systematic pipeline for that purpose, which includes the HoTMuSiC tool but also the ConSurf software [22] that computes the evolutionary conservation score of each residue, as well as annotations from UniProt [23]. This pipeline aims at selecting combinations of point mutations likely to achieve a substantial increase of the protein thermoresistance without affecting the function of the target protein. In what follows, we describe step by step how this pipeline works, as illustrated in Fig. 3:

**Figure 3.**
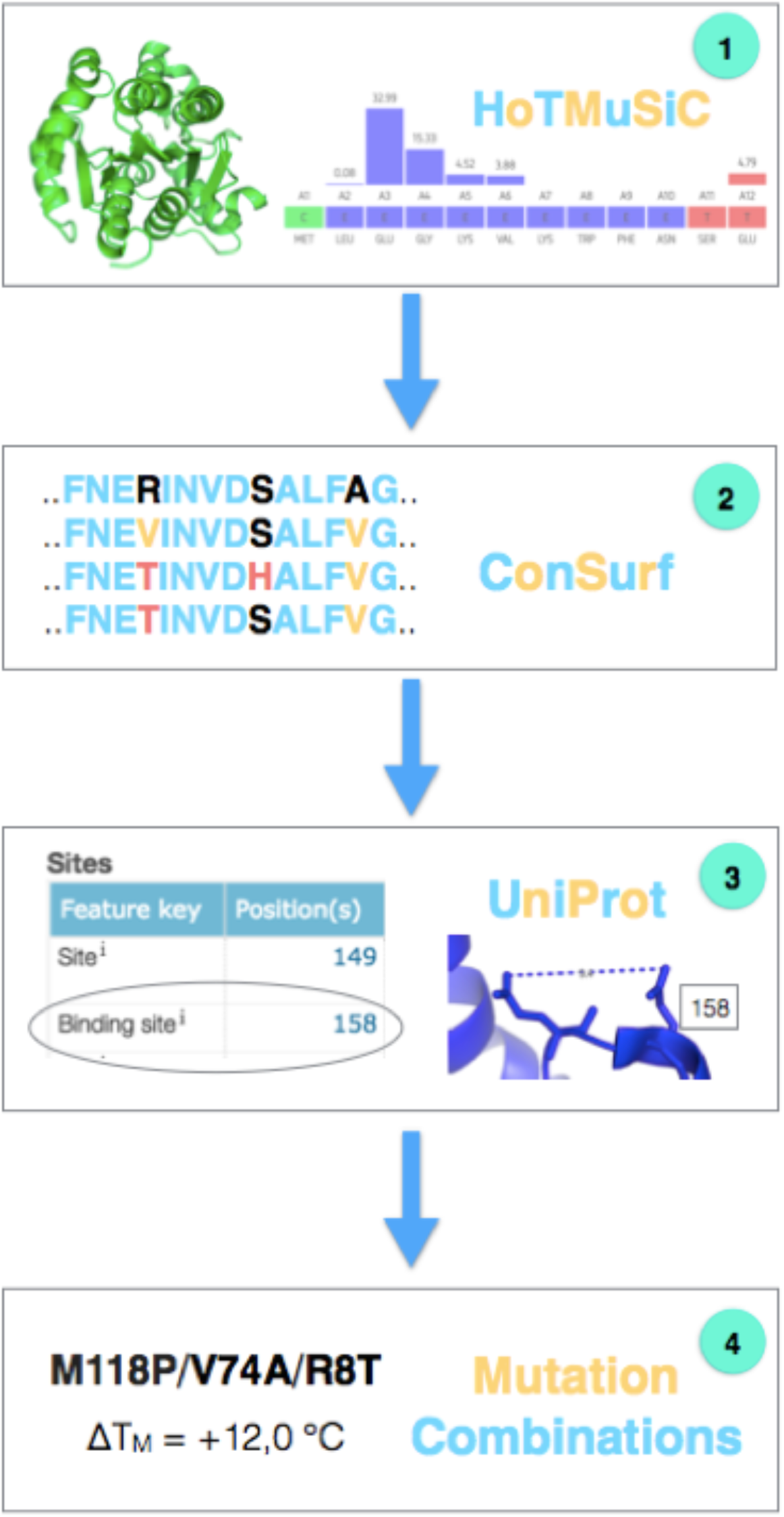
Schematic picture of the HoTMuSiC-based pipeline for protein design.

- In a first stage, the impact of all possible point substitutions on the melting temperature of a target protein is evaluated with HoTMuSiC and/or *T_m_*-HoTMuSiC in the systematic mode (see subsection 4.2), using as input the (experimental or modelled) protein structure in PDB format. If multiple PDB structures of the target protein are available, HoTMuSiC is applied to all of them (with usually a cut-off of about 2.5 Å on the X-ray resolution), and the mean Δ*T_m_* of each substitution is computed. The mutations are then classified according to their mean Δ*T_m_* values.
- The following step consists of running ConSurf [22] to estimate the evolutionary conservation score (from 0 to 9) of each amino acid of the target sequence aligned to its homologous sequences (see 3rd column of Table 3). At this level, we performed the first selection by imposing that the mutated positions have a conservation score of at most 7. This filters out mutant residues that are largely conserved and thus probably too important for functional or structural reasons.
- The next step is to retrieve from UniProt [23] the available annotations about catalytic residues and binding sites. The spatial distance between these sites and all the protein residues were computed. All positions that are closer than 7 Å (which can be computed between *C_α_* atoms, *C_β_* atoms or side chain geometric centers) from one of the functional sites were omitted. This ensures that the selected mutations do not touch or influence the catalytic activity or the binding to other biomolecules.
- The remaining list thus contains point mutations that have a high chance to improve the thermoresistance of the target protein without affecting its function. Often, however, one cannot reach the required increase in melting temperature by just a single mutation. The strategy is then to combine several point mutations. For this purpose, we made the approximation that if two mutated sites are separated by a spatial distance of more than about 10 A, the stabilization effect is additive. This final criterion leads us to select groups of two, three or more mutations that optimize the target protein.

**Table 3.** The fifteen point mutations in RML that are predicted as the most thermostabilizing by HoTMuSiC. The predicted and experimental [25] Δ*T_m_* values are reported in columns 2 and 3. The ConSurf conservation scores and the spatial distance from the catalytic residues, computed here between average side chain centroids, are shown in columns 4 and 5. The substitutions that are dropped on the basis of their conservation scores and of their distance to the catalytic triad are on a yellow and green background, respectively.

| Mutations | $\Delta T_m^{\text{pred}}$<br>(°C) | $\Delta T_m^{\text{exp}}$<br>(°C) | ConSurf | Distance<br>(Å) |
| --- | --- | --- | --- | --- |
| E221V | 3.2 | - | 9 | 6.3 |
| E221I | 2.8 | - | 9 | 6.3 |
| Q174F | 2.5 | - | 6 | 5.3 |
| Q174Y | 2.3 | - | 6 | 5.3 |
| E230F | 2.0 | 1.6 | 3 | 8.5 |
| E230I | 2.0 | 5.7 | 3 | 8.5 |
| E230Y | 1.8 | 2.1 | 3 | 8.5 |
| E230V | 1.8 | 5.4 | 3 | 8.5 |
| Q176F | 1.8 | - | 8 | 5.2 |
| E221L | 1.8 | - | 9 | 6.3 |
| Q174W | 1.8 | - | 4 | 5.3 |
| Q176Y | 1.8 | - | 8 | 5.2 |
| E230L | 1.7 | 5.4 | 3 | 8.5 |
| A64I | 1.7 | - | 8 | 8.0 |
| E230W | 1.7 | 2.7 | 3 | 8.5 |

### 4.5. HoTMuSiC-based pipeline applied to *Rhizomucor miehei* lipase

To illustrate how the HoTMuSiC pipeline can be used to optimize a protein, we applied it to thermally stabilize the lipase from *Rhizomucor miehei* (RML). This enzyme catalyses a wide range of reactions such as the hydrolysis of oil, the esterification of fatty acids, and the transesterification and alcoholysis of glycerides. Therefore, it received a lot of attention, and different studies tried to improve its thermal stability to increase the efficiency of these biocatalytic reactions [24].

There are several available X-ray structures of the enzyme, and we considered here the two structures with PDB code 3TGL and 4TGL, which have a resolution of 1.9 Å and 2.6 Å, respectively (Fig. 4). RML folds into a *β*-sheet surrounded by helices (Rossmann fold) and has a melting temperature of 58.7 °C [25]. It contains a Ser-His-Asp trypsin-like catalytic triad, in which the active serine is buried under a short helical lid that undergoes conformational changes (see Fig 4a). By exposing or protecting the catalytic pocket, the lid movement controls the enzymatic activity [25].

**Figure 4.**
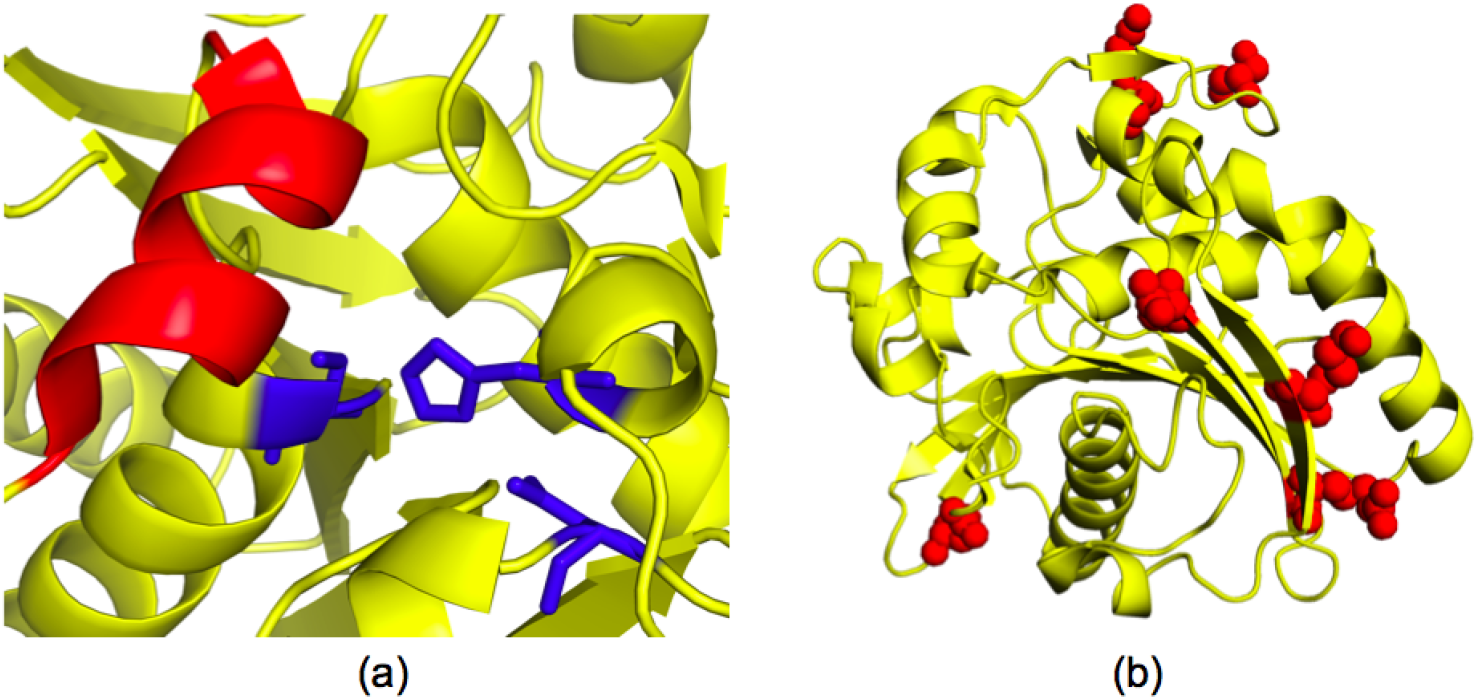
3D structure of Rhizomucor miehei lipase (PDB code 3TGL) (a) Zoom on the Ser-His-Asp catalytic triad (in blue sticks) with the helical lid (in red) that undergoes conformational changes that modulate the protein activity. (b) Residues to be mutated (in red spheres) for the thermostabilization of the lipase.

The first step to predict thermostabilizing mutations consists in running HoTMuSiC in the systematic mode on the two PDB structures of RML to compute the Δ*T_m_* of all possible point mutations. These values, stored in the 3TGL.hot and 4TGL.hot files on the Results page, were first averaged for each individual mutation and then ranked decreasingly according to their average Δ*T_m_* value.

The top fifteen mutations in the list are given in Table 3. Note that glycine and proline substitutions were excluded from this list since their mutations are likely to induce local changes in conformation which are not taken into account in the prediction model.

In the next steps, the residue conservation across homologous sequences was evaluated using the ConSurf algorithm [22], and the spatial distance between the mutated residues and the catalytic triad Ser-His-Asp was computed. The mutations inserted at highly conserved positions (ConSurf score ≥ 8) or that are too close to the catalytic site (distance ≤ 7Å) are then removed, as they might impact the protein function or other biophysical characteristics that we do not want to modify.

The substitutions filtered out due to their high conservation scores or their proximity to the catalytic triad, and those that satisfy all selection criteria, are shown in Table 3. Among the 15 top mutations predicted in step 1, we kept only the six mutations of Glu at position 230.

To get a larger number of candidate mutations, we have to relax the first, Δ*T_m_*-based, criterion. In this way we obtained the fifteen most stabilizing mutations that satisfy all our selection filters, shown in Table 4 and Fig. 4b.

**Table 4.**
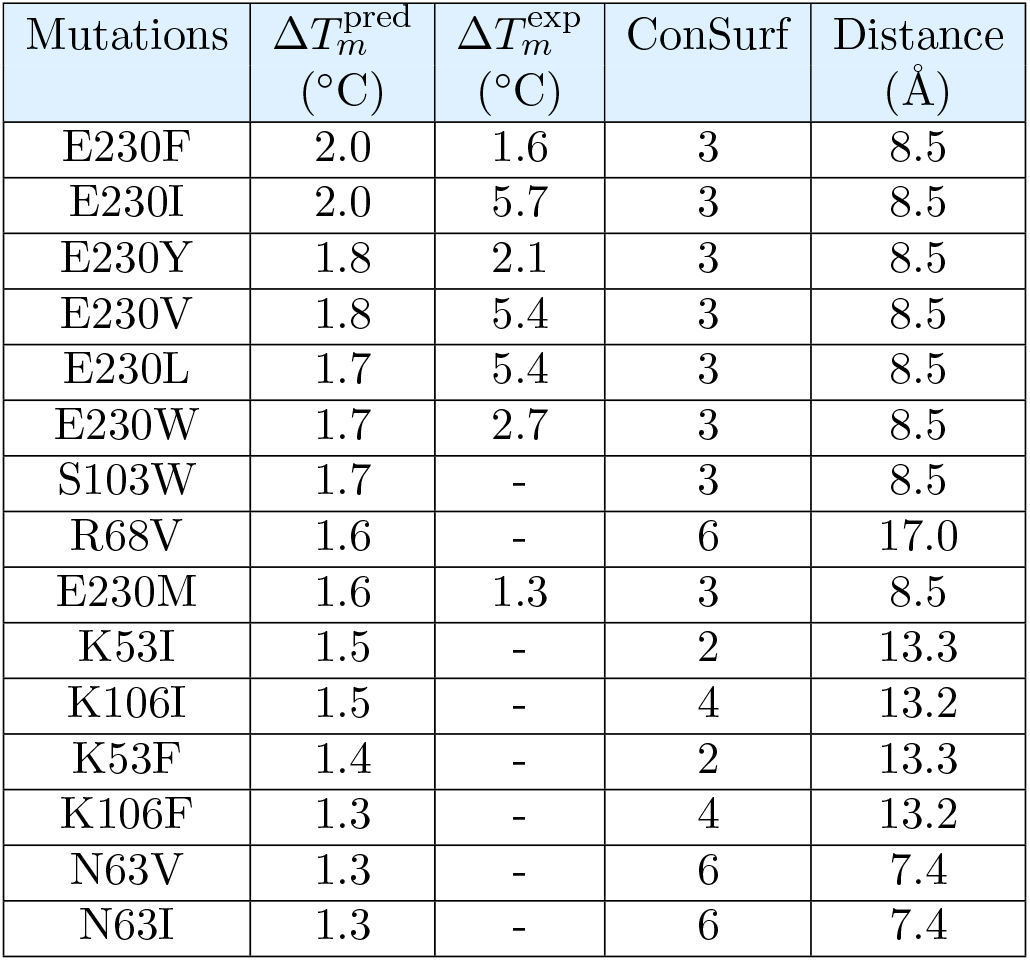
Final list of proposed mutations in RML, which satisfy all our selection criteria.

As a final step of our pipeline, we determined subsets of mutations of residues whose relative distances are equal to 10 Å or more, and are thus assumed to be independent. For example, this procedure yields the sets of multiple mutations:

- E230I/S103W/K53I
- E230F/R68V/K53F
- E230M/K106I/N63V

Finally, we compared our computational predictions on lipase with the available experimental data reported in [25] and shown in Tables 3 and 4. The prediction score of HoTMuSiC on this set of 36 point mutations was found to outperform the competitor methods reported in Table 1 and to reach quite a good accuracy evaluated by a root mean square deviation between predicted and experimental Δ*T_m_* values of 1.7 °C and by a linear correlation coefficient of 0.6. This score is the same as the one obtained from the other tests of HoTMuSiC shown in Tables 1-2. Moreover, four sets of multiple mutations have been experimentally shown to lead to a substantial increase of the RML thermoresistance [25]. If we assume that the effect of mutations with a sufficient spatial separation is additive, these multiple mutations are also very well predicted by HoTMuSiC. The final score of our pipeline, when the four multiple mutations are added to the 36 point mutations, reaches a linear correlation coefficient higher than 0.8.

Finally note that the predicted Δ*T_m_* values tend to be lower than the experimental values. Indeed, as we already noted [26], the training datasets are dominated by destabilizing mutations and this induces biases toward these types of mutations and leads to an underestimation of the predicted stabilization effects.

## 5. Conclusion

In this chapter, we presented recently developed protein thermal stability predictors, and their application to efficiently optimize targeted proteins. We focused more specifically on the HoTMuSiC predictor, which predicts the Δ*T_m_* values of all the possible point mutations in a medium-size protein in a few minutes, on the basis of its experimental or modelled 3D structure. Our tool outperforms the other available Δ*T_m_* predictors, as well as ΔΔ*G*-predictors when applied to thermal stability even though they are designed for thermodynamic stability.

The fastness of HoTMuSiC allows scanning and predicting all possible substitutions inserted in a target protein. It can thus guide mutagenesis experiments aimed to improve the thermoresistance of proteins, and be employed in the optimization of a wide series of biotechnological processes.

To optimize the selection of mutations that need to be tested experimentally, we complemented the HoTMuSiC results with additional information on the residue conservation among homologous proteins and the distance from annotated functional sites. This novel pipeline is designed to filter out mutations that are likely to affect functionally or structurally important residues. The point mutations that satisfy the criteria are then combined into subsets of non-interacting mutations that are assumed to be independent. These subsets are identified to strongly enhance the effect on thermal stability.

The HoTMuSiC pipeline was applied to the thermal stabilization of lipase from *Rhizomucor miehei*. Several series of multiple mutations were predicted for this enzyme. For those mutations whose Δ*T_m_* values were measured experimentally, the prediction score was shown to be high. Finally note that the potentiality of HoTMuSiC is not restricted to the protein design field. Due to its speed, it can also be applied on a proteomic scale to gain important theoretical insights into the thermal and evolutionary adaptation of proteins to extreme environments.

## 6. Acknowledgement

We thank Raphael Bourgeas for the help in implementing HoTMuSiC on the webserver, and Cesar Ngabo for his contribution to the HoTMuSiC pipeline. We acknowledge support from the Fund for Scientific Research - FNRS through a PDR research project. FP and MR are FNRS postdoctoral researcher and research director, respectively.

